# EMX2 transcriptionally regulates *Nfib* expression in neural progenitor cells during early cortical development

**DOI:** 10.1101/2021.12.26.474186

**Authors:** Jonathan W. C. Lim, Jens Bunt, Caitlin R. Bridges, Ching Moey, Matisse T. Jacobs, Kok-Siong Chen, Linda J. Richards

**Affiliations:** The University of Queensland, Queensland Brain Institute, Brisbane, Queensland, Australia; The University of Queensland, School of Biomedical Sciences, Brisbane, Queensland, Australia

**Keywords:** cortical development, EMX2, neuronal differentiation, nuclear factor one, NFIB

## Abstract

The nuclear factor one (NFI) transcription factors play key roles in regulating the onset of both neuronal and glial differentiation during cortical development. Reduced NFI expression results in delayed differentiation, which is associated with neurodevelopmental disorders in humans that include intellectual disability, agenesis of the corpus callosum and macrocephaly. Despite their importance, our understanding of how individual NFI family members are regulated during cortical development remains limited. Here, we demonstrate that in mice, the homeobox transcription factor EMX2 regulates *Nfib* expression in radial glial cells during cortical development. Using a combination of bioinformatics, molecular and histological approaches, we demonstrate that EMX2 is able to bind to the *Nfib* promoter to up-regulate *Nfib* expression. Unexpectedly, *in vivo* over-expression of EMX2 in wildtype animals does not further up-regulate NFIB but instead leads to its down-regulation. Therefore, our findings suggest that EMX2 is capable of both activating and repressing *Nfib*, in a context-dependent manner. This bi-directional control over *Nfib* expression enables fine-tuning of the total level of NFI proteins expressed and could be important for cell-type specific NFI functions.

## Introduction

During development, the timely onset of cellular differentiation is essential to generate the basic anatomic architecture that is required for a fully functional brain. Underpinning this process is the regulation of gene expression, governed by transcription factors that are dynamically expressed in precise spatiotemporal patterns. One such family of transcription factors that are required for the timely differentiation of neural progenitors are the nuclear factor one (NFI) transcription factors. Analyses in mice demonstrate that the expression of these transcription factors in the developing cortex is first detected at approximately embryonic day (E)11.5 (Chaudhry et al. 1997; Plachez et al. 2008), coinciding with the onset of neurogenesis within this region. In humans, haploinsufficiency of the genes *NFIA*, *NFIB* or *NFIX* result in a spectrum of syndromes that are characterised by neurodevelopmental deficits that include macrocephaly, other varying brain malformations and intellectual disability (Zenker et al. 2019).

*Nfi* knockout mouse models recapitulate the phenotypes reported in humans. These are grossly represented by an expansion of the lateral ventricles and dysgenesis of the corpus callosum, both of which arise due to delayed differentiation of radial glial cells (das Neves et al. 1999; Shu et al. 2003; Barry et al. 2008; Campbell et al. 2008; Piper et al. 2009; Betancourt et al. 2014; Gobius et al. 2016). During embryonic development, the delay in radial glial cell differentiation in the developing cortex is accompanied by the continued self-renewal of these cells. Consequently, the ventricular zone (VZ) where radial glial cells reside expands and the cortical plate is thinner. Analyses of mouse models at postnatal ages demonstrate enlarged cerebral cortices as compared to wildtype littermates (Campbell et al., 2008; Schanze et al. 2018). This enlargement is presumably due to the subsequent differentiation of the expanded progenitor cell pool that eventually occurs in these mice. As evident by the increased severity of the phenotype when multiple family members are concurrently deleted (Harris et al. 2016; Bunt et al. 2017), tight control of NFI levels is requisite for normal development. Despite their importance, little is known about how the expression of the NFI transcription factors is regulated during brain development.

The transcription factor EMX2 is expressed from approximately E8 in telencephalic progenitors within the dorsal and medial VZ (Simeone et al. 1992; Gulisano et al. 1996; Mallamaci et al. 1998). Analyses of knockout mouse models demonstrate that EMX2 expression during this early period is critical for the specification of the medial cortex. In the absence of EMX2, the hippocampus and cingulate cortex are considerably smaller because of precocious differentiation and depletion of the progenitor cell population (Pellegrini et al. 1996; Yoshida et al. 1997; Shinozaki et al. 2002; Muzio et al. 2005). The neocortical phenotype of *Emx2* knockout mice is considerably milder. Nevertheless, EMX2 plays complex autonomous and non-cell autonomous roles in this region. Among its known roles are regulating the maintenance of the progenitor cell population (Heins et al. 2001; Brancaccio et al. 2010), the differentiation and migration of specific neuronal populations (Mallamaci et al. 2000; Shinozaki et al. 2002), and cortical arealisation (Yoshida et al. 1997; Mallamaci et al. 2000; Bishop et al. 2000; Bishop et al. 2002). Therefore, while EMX2 expression is restricted to progenitor cells occupying the VZ, its expression in these cells is critical for regulating many cellular processes throughout cortical development.

Here, we utilised a bioinformatics approach to identify EMX2 as a candidate transcriptional regulator of *Nfib*. Using a combination of histological and molecular analyses, we further demonstrate that EMX2 is required to drive NFIB expression in radial glial cells during cortical development. Unexpectedly, ectopic over-expression of EMX2 in the developing cortex does not further up-regulate NFIB but instead leads to its down-regulation. Therefore, our findings suggest that EMX2 could function by fine-tuning NFIB expression and is capable of both activating and repressing *Nfib* in this regard.

## Material and Methods

### Animal breeding and tissue collection

All breeding and experiments were performed at the University of Queensland in accordance with the Australian Code of Practice for the Care and Use of Animals for Scientific Purposes and with approval from the University of Queensland Animal Ethics Committee. *Nfib* knockout (*Nfib^tm1Rmg^*) (Steele-Perkins et al. 2005) and *Emx2* knockout (*Emx2^tm1Pgr^*) (Pellegrini et al. 1996) mice were maintained on a C57Bl/6 background. To generate timed-pregnant females, male and female mice were placed together overnight and checked the following day for vaginal plugs. This day was designated as E0.5 if a vaginal plug was present. Pregnant dams were euthanised using sodium pentobarbital (Abbott Laboratories) on the day of embryo collection. E13.5 embryos were drop fixed in 4% (w/v) paraformaldehyde (PFA) in phosphate-buffered saline (PBS; pH 7.4) for fluorescence immunohistochemistry or their neocortex dissected in ice-cold sterile PBS and immediately snap frozen for mRNA isolation. E15.5 embryos were transcardially perfused with 0.9% (w/v) saline followed by 4% PFA.

### Bioinformatics analyses

Candidate transcriptional regulators of *Nfib* were identified using FIMO software version 5.2.0 (Bailey et al. 2009; Grant et al. 2011). Briefly, the region encompassing the mouse *Nfib* RefSeq transcription start site and 2000 base pairs upstream and downstream of this (chr4:82503780-82507779 from mm10 assembly) was scanned for putative transcription factor binding sites (p < 0.0001) obtained from a database of transcription factor motifs (Jolma et al. 2013). Candidate transcriptional regulators were then filtered to remove transcription factors with low or undetectable expression in the E11.5 forebrain based on expression data from ENCODE mRNA-seq datasets (transcripts per million < 10; dataset identifiers: ENCFF465SNB, ENCFF976OLT) (ENCODE Project Consortium 2012; Davis et al. 2018; He et al. 2020) and the Allen Developing Mouse Brain Atlas (http://developingmouse.brain-map.org) (Thompson et al. 2014). Pairwise alignment of the mouse and human *Nfib* promoter was performed using the EMBOSS Water webtool (Madeira et al. 2019). A subsequent scan for EMX2 binding sites using a less stringent significance threshold (p < 0.001) was then performed using FIMO. mRNA expression data of 6 to 16 postconceptional weeks (PCW) human cortical tissue from the BrainSpan Transcriptional Atlas of the Developing Human Brain (http://brainspan.org) (Miller et al. 2014) and GSE25219 (Kang et al. 2011) datasets were analysed and visualised using the R2: Genomics Analysis and Visualization Platform (http://r2.amc.nl) as previously described (Bunt et al. 2010; Bunt et al. 2012). Single-cell mRNA-seq data of SOX2+/PAX6+ cells from 14 to 19 PCW human cortical samples (Thomsen et al. 2016) and E14.5 mouse cortical radial glial cells (Loo et al. 2019) were tested for statistical significance using Pearson’s correlation test in Prism 7 (GraphPad Software).

### Immunohistochemistry

Brains of PFA-fixed embryos were removed from the skull and sectioned coronally, sagittally or horizontally at 50 μm on a vibratome. Fluorescence immunohistochemistry was conducted as previously described with minor modifications (Plachez et al. 2008). The primary antibodies that were used are: rabbit anti-NFIB (1:500; HPA003956, Atlas Antibodies), chicken anti-β-galactosidase (1:500; ab9361, Abcam), chicken anti-GFP (1:1000; ab13970, Abcam) and rabbit anti-EMX2 (1:500; HPA065294, Atlas Antibodies). Validation of the specificity of the EMX2 antibody is presented in Supplementary Fig. 1. The specificity of the NFIB antibody was validated previously (Chen et al. 2017). Alexa Fluor 488-conjugated goat anti-chicken (1:500; A-11039, Invitrogen) and Alexa Fluor 555-conjugated goat anti-rabbit (1:500; A-11034, Invitrogen) secondary antibodies were used for detection.

Immunofluorescent sections were counterstained with 4’,6-diamidino-2-phenylindole, dihydrochloride (DAPI, Invitrogen) to label cell nuclei and coverslipped using ProLong Gold anti-fade reagent (Invitrogen). To reduce technical variability when quantifying immunofluorescence intensity, matched control sections were mounted on the same slide to ensure identical staining conditions.

### Imaging and data analysis

High resolution fluorescence images were acquired using a Diskovery spinning disk confocal system (Spectral Applied Research, Ontaria, Canada) built around a Nikon TiE body equipped with two sCMOS cameras (Andor Zyla 4.2, 2048 x 2048 pixels) and captured with Nikon NIS software (Nikon, Tokyo, Japan). For fluorescence intensity measurements across the different cortical layers, matched neocortical regions of 100 μm width were cropped and analysed using ImageJ software (NIH). To generate fluorescence intensity histograms, cropped sections were divided into 20 bins of equal size across the ventricular to pial surfaces. Fluorescence intensity measurements for *in utero* electroporations were determined by measuring the mean grey value of individual nuclei within the electroporated region. Statistical significance was determined using unpaired t-tests unless otherwise stated, with p-values below 0.05 considered significant. All values are presented as the mean unless otherwise stated, with error bars representing the standard error of the mean. Images in figures are representative images that have been identically cropped, enhanced for contrast and brightness, and pseudo-colored to permit overlay using Adobe Photoshop software.

### RNA isolation and quantitative PCR

Total RNA was extracted from E13.5 snap frozen neocortical tissue using TRIzol reagent (Invitrogen) as per the manufacturer’s protocol (Life Technologies, USA). RNA was reverse transcribed using SuperScript III First-Strand Synthesis SuperMix (Invitrogen) and Oligo(dT) primers (Invitrogen) as per the manufacturer’s instructions. Real-time qPCR was performed with Platinum SYBR Green qPCR SuperMix-UDG (Invitrogen) and 0.25 μM forward and reverse primers on a Rotor-Gene 3000 (Corbett Life Science) as previously described (Bunt et al. 2015). Relative expression was determined using the ΔΔCt method with the housekeeping gene beta-2-microglobulin used as a relative standard. Statistical significance was determined using Welch’s t-test. Error bars represent the standard error of the mean. Primers were as previously described (Messina et al. 2010): B2MM 5’-agactgatacatacgcctgcag-3’, B2MMC 5’-gcaggttcaaatgaatcttcag-3’, mNFI-BE2 5’-gtttttggcatactacgtgcagg-3’ and mNFI-BE3C 5’-ctctgatacattgaagactccg-3’.

### Plasmid DNA

The mouse *Emx2* coding sequence was amplified from pMXIG-Emx2 (a kind gift from Magdalena Götz) using the following primers containing EcoRI restriction sites: 5’-ATCGGAATCATGTTTCAGCCGGCGCCCAAG-3’ and 5’-ATCGGAATTCTTAATCGTCTGAGGTCACATC-3’. The amplified fragment was subsequently digested and cloned into the EcoRI site within pCAGIG (Addgene plasmid #11159; a kind gift from Connie Cepko) (Matsuda and Cepko 2004) to generate pCAGIG-Emx2. Cloning of the pNfib luciferase plasmid was previously described (Bunt et al. 2015).

### Dual-luciferase reporter assays

Human glioblastoma U-251 MG (Ponten and Macintyre 1968) and mouse neuroepithelial NE-4C (Schlett and Madarasz 1997) cell lines were obtained from the American Type Culture Collection (ATCC). Cells were cultured in DMEM (Invitrogen) supplemented with 10% fetal bovine serum at 37°C in a humified atmosphere containing 5% CO_2_. Cells were seeded for luciferase reporter assays into a 48-well (U-251) or 96-well (NE-4C) plate 24 hours prior to transfection. Either pCAGIG-Emx2 or pCAGIG were cotransfected with pNfib or the control pGL4.23 luciferase reporter plasmid (Promega) using FuGENE HD (Promega). A control *Renilla* luciferase plasmid (pRL-SV40; Promega) was co-transfected with all transfections as an internal control for normalization of firefly luciferase activity. Luciferase activity was assayed 48 hours after transfection using the Dual-Luciferase Reporter Assay System (Promega). All experimental conditions were tested as three or four independent experiments consisting of technical triplicates. Statistical significance was determined for each cell line using ratio paired t-tests.

### ChIP-qPCR

NE-4C cells were cultured in DMEM (Invitrogen) supplemented with 10% fetal bovine serum at 37°C in a humified atmosphere containing 5% CO_2_. 100 mm cell culture dishes were seeded with cells 24 hours prior to transfection and transfected the following day with the pCAGIG-Emx2 plasmid using FuGENE 6 transfection reagent (Promega) as per the manufacturer’s recommendations. Cells were fixed 24 hours after transfection using 1% (w/v) PFA in PBS for 5 minutes. Excess PFA was then quenched with 125 mm glycine. Fixed cells were lysed in lysis buffer (10 mM Tris-HCl pH 8.1, 10 mM NaCl, 3 mM MgCl_2_, 0.5% (v/v) Nonidet P40 substitute solution) supplemented with cOmplete protease inhibitor cocktail (Roche) at 4°C for 20 minutes in order to isolate fixed nuclei. Isolated nuclei were sonicated in sonication buffer (0.1% (w/v) SDS, 1 mM EDTA pH 8.0, 10 mM Tris-HCl pH 8.1) supplemented with cOmplete protease inhibitor cocktail (Roche) using the M220 focused-ultrasonicator (Covaris) to generate 200 to 500 base pairs chromatin fragments. Chromatin immunoprecipitation (ChIP) was performed by combining chromatin fragments isolated from 2.5 × 10^6 cells and suspended in ChIP buffer (0.087% (w/v) SDS, 0.87 mM EDTA pH 8.0, 8.7 mM Tris-HCl pH 8.1, 150 mM NaCl, 1% (v/v) Triton X-100) supplemented with cOmplete protease inhibitor cocktail (Roche) with 5 μg primary antibody and 50 uL Pierce Protein G magnetic beads (Thermo Scientific). The primary antibodies used were rabbit anti-EMX2 (HPA065294, Atlas Antibodies) and normal rabbit IgG (#2729, Cell Signaling Technology). Following ChIP, chromatin-bound beads were washed twice with low salt wash buffer (0.1% (w/v) SDS, 1% (v/v) Triton X-100, 2mM EDTA pH 8.0, 150 mM NaCl, 20 mM Tris-HCl pH 8.1), high salt wash buffer (0.1% (w/v) SDS, 1% (v/v) Triton X-100, 2mM EDTA pH 8.0, 500 mM NaCl, 20 mM Tris-HCl pH 8.1), LiCl wash buffer (1% (w/v) sodium deoxycholate, 0.1% (v/v) Nonidet P40 substitute solution, 2 mM EDTA pH 8.0, 250 mM LiCl, 10 mM Tris-HCl pH 8.1) and TE buffer (10 mM Tris-HCl pH 8.1, 1 mM EDTA pH 8.0). Chromatin was eluted from beads using 0.1 M sodium bicarbonate and 1% (w/v) SDS at 65°C for 20 minutes. Eluted chromatin was de-crosslinked at 65°C for 20 hours and sequentially treated with 0.2 mg/mL RNase A (37°C for 1 hour) and 0.4 mg/mL Proteinase K (55°C for 1 hour) before being purified using the QIAquick PCR purification kit (QIAGEN). qPCR was performed on the Rotor-Gene 3000 (Corbett Life Science) using the PowerUp SYBR Green Master Mix (Life Technologies) and the following thermocycler conditions: 2 minutes at 50°C and 2 minutes at 95°C, followed by 40 cycles with 15 seconds denaturation at 95°C and 1 minute annealing and extension at 60°C. Relative enrichment is presented at percent input and statistical significance was determine using unpaired t-tests. Primers used for ChIP-qPCR were: site 1 forward 5’-gagaggctggtgcaaaagg-3’, site 1 reverse 5’-ttgaaagattcaccggttcc-3’, site 2 forward 5’-tttaaccccgtcctgtcctc-3’, site 2 reverse 5’-aggctcttggtttacagggg-3’, downstream control forward 5’-actttaggcaccccaaaacg-3’, downstream control reverse 5’-gggagcgtctgtagatggac-3’, upstream control forward 5’-acccagagagcccaagattc-3’ and upstream control reverse 5’-catagccaccatcagcactg-3’. Statistical significance was determined using unpaired t-tests.

### In utero *electroporation*

*In utero* electroporation was performed as previously described (Lim et al. 2015). Briefly, pregnant wildtype mice on a CD1 background were anaesthetised with 80-100 mg/kg (body weight) ketamine and 5-10 mg/kg xylazine. An abdominal incision was made to expose the uterus, and 0.5-1.0 μl plasmid DNA (pCAGIG or pCAGIG-Emx2) diluted in sterile PBS was injected through the uterine wall into the medial region of the lateral ventricle of the brain with a pulled glass pipette attached to a Picospritzer III (Parker Hannifin). Electrical pulses (five 35 V pulses of 50 ms applied at 1 second intervals) were then delivered to embryos by electrodes connected to a square-wave ECM 830 electroporator (BTX Harvard Apparatus). Embryos were returned into the abdominal cavity and the abdominal cavity was sutured. Isotonic Ringer’s solution was administered subcutaneously and dams were placed in a heated recovery chamber until alert. Buprenorphine (0.05-0.15 mg/kg) was administered orally as an analgesic through ingestion of a flavoured jelly injected with buprenorphine.

## Results

### EMX2 is a candidate transcriptional regulator of *Nfib* expression

*Nfib* expression in the developing cortex is first observed at approximately E11.5 (Chaudhry et al. 1997; Plachez et al. 2008) when the cortex is composed primarily of radial glial cells and neurogenesis is just beginning. To identify transcriptional regulators of *Nfib* expression, we undertook a bioinformatics approach by identifying transcription factor motifs present within the *Nfib* promoter and subsequently filtering candidate transcription factors based on their expression at E11.5 in ENCODE mRNA-seq data (Thompson et al. 2014) and the Allen Developing Mouse Brain Atlas (He et al. 2020). Using this approach, we identified EMX2 as a potential regulator of *Nfib* expression. Putative EMX2 binding sites were present within the promoter regions of both human and mouse *Nfib* (Fig. 1A). These binding sites occurred within two highly conserved regions that potentially function as transcriptional regulatory elements (grey boxes in Fig. 1A). Besides these, no further EMX2 binding sites were observed within 2000 base pairs upstream or downstream of the *Nfib* transcriptional start site.

**Fig. 1.**
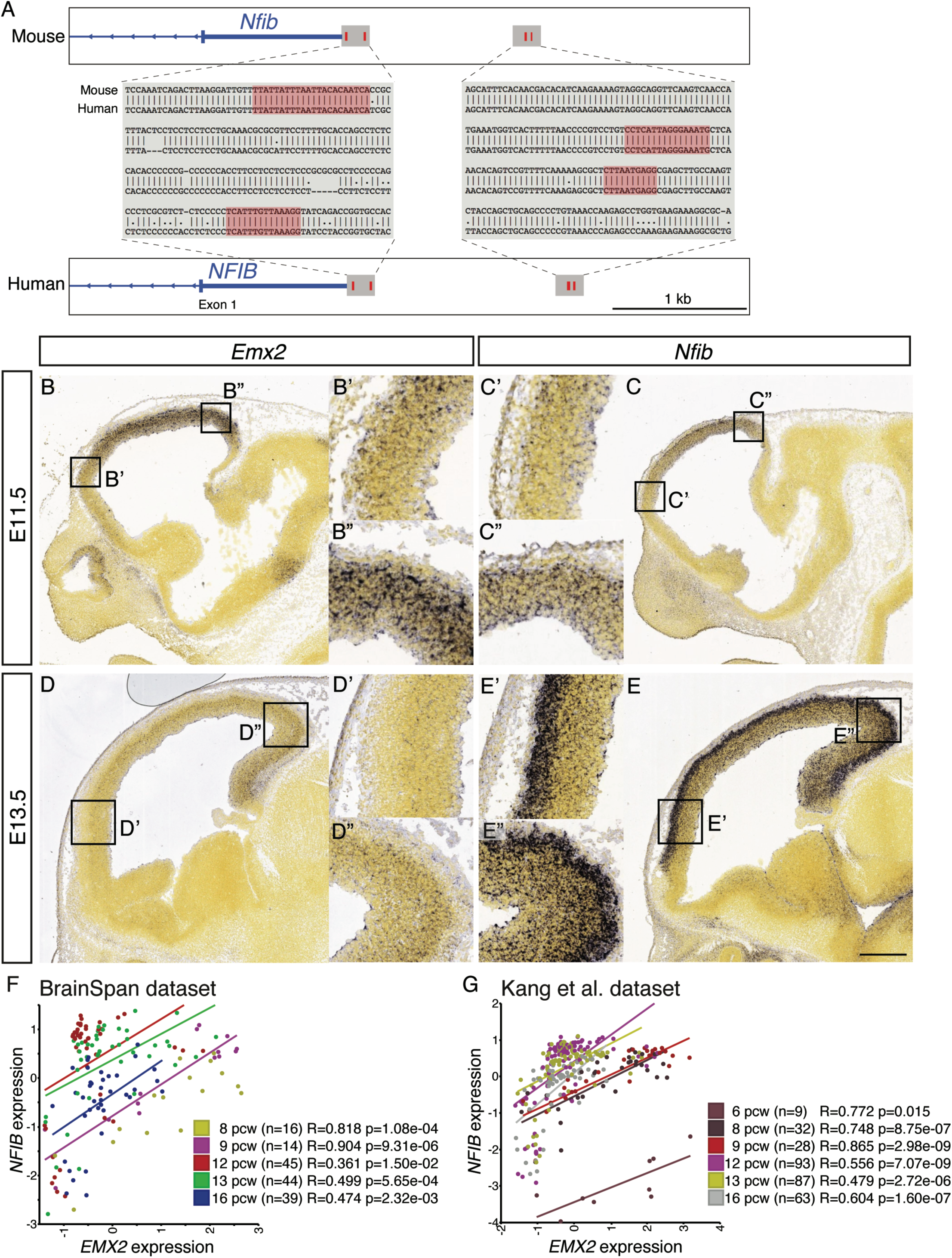
EMX2 is a candidate transcriptional regulator of NFIB. (*A*) Schematic of the proximal promoter region of *Nfib* in mouse and human. Solid blue bars represent the 5’ untranslated region and exon 1 of *Nfib*. Red boxes denote potential EMX2 binding sites as identified using the motif finding tool FIMO (Grant et al. 2011), with the grey boxes demonstrating sequence conservation within these regions (mouse mm10 and human hg38). Scale bar represents 1000 base pairs. (*B*-*E*) *In situ* hybridization images of E11.5 and E13.5 sagittal sections probed for *Emx2* and *Nfib* expression were obtained from the Allen Developing Mouse Brain Atlas (Thompson et al. 2014), demonstrating similar expression gradients. Scale bar represents 400 μm. Image credit: Allen Developing Mouse Brain Atlas (http://developingmouse.brain-map.org). Image identifiers: image 7 of 18 from experiment 100047257 (E11.5 *Emx2*), image 11 of 16 from experiment 100041799 (E13.5 *Emx2*), image 7 of 16 from experiment 100054279 (E11.5 *Nfib*), and image 14 of 18 from experiment 100054417 (E13.5 *Nfib*). (*F*, *G*) Correlation of *EMX2* (x-axis) and *NFIB* (y-axis) mRNA expression in human fetal brain samples from 6 to 16 postconceptional weeks (PCW) from Brainspan (Miller et al. 2014) and GSE25219 (Kang et al. 2011), respectively. Each colour represents a different PCW.

EMX2 expression during cortical development is detected as early as E8.5, where it is expressed in neuroepithelial cells and subsequently in radial glial cells in high caudal to low rostral and high medial to low lateral gradients (Simeone et al. 1992; Gulisano et al. 1996; Mallamaci et al. 1998). While NFIB expression is not limited to the VZ during cortical development, we previously reported that NFIB expression in radial glial cells followed a similar expression pattern (Bunt et al. 2015). These patterns of expression are replicated within *in situ* hybridization data from the Allen Developing Brain Atlas (Fig. 1B-E). In E11.5 wildtype sagittal sections, the highest expression of *Emx2* and *Nfib* mRNA were observed in the hippocampal primordium and the caudal region of the dorsal cortex, with lower expression detected in rostral regions (Fig. 1B, C). This expression pattern was similarly observed at E13.5 in cells occupying the germinal layer but not within the nascent cortical plate (Fig. 1D, E). At this age, the nascent cortical plate strongly expresses *Nfib*, but *Emx2* is absent within this layer. Furthermore, *Nfib* expression within the cortical plate appears more evenly distributed and does not conform to the graded expression pattern observed within the VZ. These findings suggest that EMX2 could be a transcriptional regulator of *Nfib* within the VZ but is not required to sustain *Nfib* expression in differentiated neurons.

Given that potential EMX2 binding sites identified in the mouse *Nfib* promoter are conserved within the human *NFIB* promoter, we extended our analyses using human RNA-seq data to determine whether the relationship between *Emx2* and *Nfib* expression was similarly conserved in humans. To do this, we first analysed the expression of *EMX2* and *NFIB* mRNA in two independent spatio-temporal transcriptome data sets of human foetal brain samples collected between 6 and 16 PCW (approximately equivalent to E10 to E15.5 in mice) (Kang et al. 2011; Miller et al. 2014). *NFIB* mRNA expression was similarly correlated with *EMX2* expression in both these data sets (Fig. 1F, G). We also extended our analyses to single-cell RNA sequencing (scRNA-seq) data that was generated from neural progenitor cells of the human and mouse cortex. In a dataset consisting of 186 SOX2^+^, PAX6^+^ cells isolated from 14 to 19 PCW human cortical tissues (Thomsen et al. 2016), *NFIB* expression was significantly correlated with *EMX2* expression (Pearson’s correlation: r= 0.3086, p<0.0001). Similarly, mouse *Nfib* expression is correlated with *Emx2* expression in cortical radial glial cells isolated from the E14.5 mouse cortex (Pearson’s correlation: r= 0.3782, p<0.0001; n = 1605 cells classified as radial glial cells post hoc via unsupervised cell clustering) (Loo et al. 2019). Hence, EMX2 could function as a transcriptional activator of *NFIB* expression during early cortical development in both humans and mice.

### EMX2 and NFIB proteins are co-expressed within the ventricular zone during cortical development

We further characterised the expression patterns of EMX2 and NFIB protein by immunofluorescence analyses. To do this, we examined co-expression of EMX2 and the β-galactosidase (βGal) reporter protein in E13.5 *Nfib* heterozygous brains sectioned sagittally, horizontally and coronally. The βGal gene acts as a reporter of NFIB expression as it replaces the second exon of the *Nfib* null allele and is driven by the *Nfib* promoter. Therefore, βGal expression recapitulates expression of endogenous NFIB protein (Steele-Perkins et al. 2005; Piper et al. 2009; Betancourt et al. 2014; Bunt et al. 2015). Immunofluorescence analyses demonstrate that EMX2 and βGal were both highly expressed within the developing dorsal telencephalon at E13.5 (Fig. 2). Lower expression of both proteins was detected in other brain regions and in these regions were predominantly confined to the lining of the ventricles (Fig. 2A). Within the dorsal telencephalon, the expression of EMX2 and βGal was identical to that observed for *Emx2* and *Nfib* mRNA via *in situ* hybridization. Both proteins were co-expressed within the cortical VZ in high caudal to low rostral and high medial to low lateral gradients (Fig. 2B-F). Similarly, EMX2 was not expressed within the nascent cortical plate and βGal staining appeared more evenly distributed within this region.

**Fig. 2.**
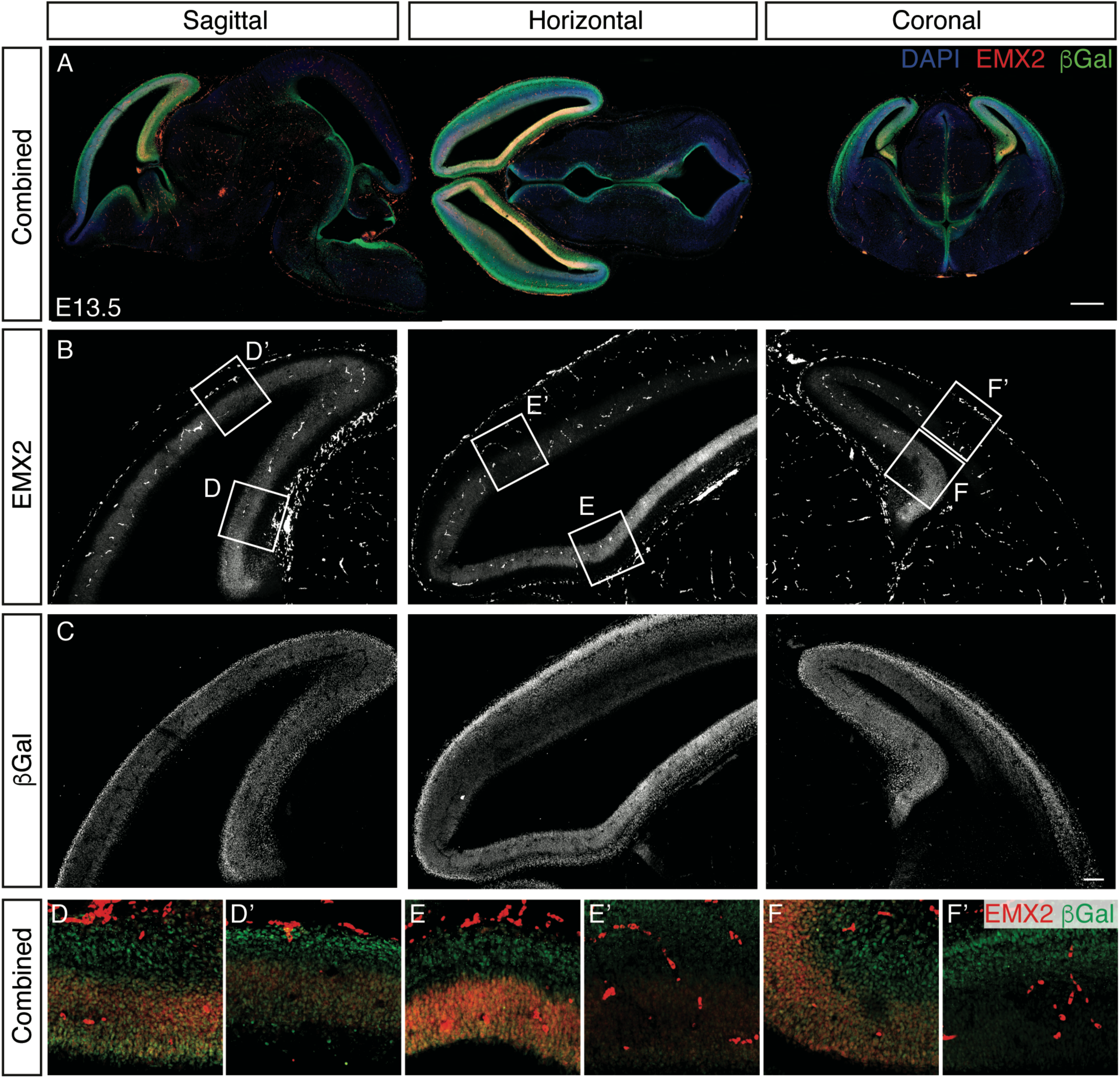
Similar expression patterns of EMX2 and NFIB in the developing brain. (*A*) Representative immunofluorescent images of sagittal, horizontal and coronal sections of E13.5 *Nfib* heterozygous brains stained for EMX2 (red) and a β-galactoside (βGal) reporter driven through the *Nfib* locus (green). Scale bar represents 500 μm. (*B*, *C*) High-powered monochrome images of the dorsal telencephalon stained for EMX2 and βGal, respectively. Scale bar represents 100 μm. (*D*-*F*) High-powered immunofluorescent images of different brain regions represented by insets in (*B*).

### Deletion of *Emx2* reduces NFIB expression during early development

To further examine the relationship between EMX2 and NFIB expression, we next investigated whether NFIB expression in the developing cortex is altered upon *Emx2* deletion. To do this, we first quantified *Nfib* expression in *Emx2* knockout and wildtype neocortical tissue by qPCR. Relative *Nfib* expression in *Emx2* knockout embryos was reduced by half as compared to wildtype littermates at E13.5 (Fig. 3A). We also quantified *Nfib* expression in these mice at E15.5 but observed no significant differences at this age (data not shown). This discrepancy between the ages could potentially be attributed to differences in the cellular composition of *Emx2* knockout tissue samples collected at different ages (Pellegrini et al. 1996; Shinozaki et al. 2002). Furthermore, while the expression of EMX2 is confined to radial glial cells that reside within the cortical VZ (Gulisano et al. 1996; Mallamaci et al. 1998), NFIB is expressed not only in radial glial cells but also in postmitotic neurons (Plachez et al. 2008; Piper et al. 2009; Betancourt et al. 2014; Bunt et al. 2015). Hence, if EMX2 regulates *Nfib* in a cell autonomous manner, the reduction in *Nfib* expression is likely to be restricted to radial glial cells that co-express both EMX2 and NFIB.

**Fig. 3.**
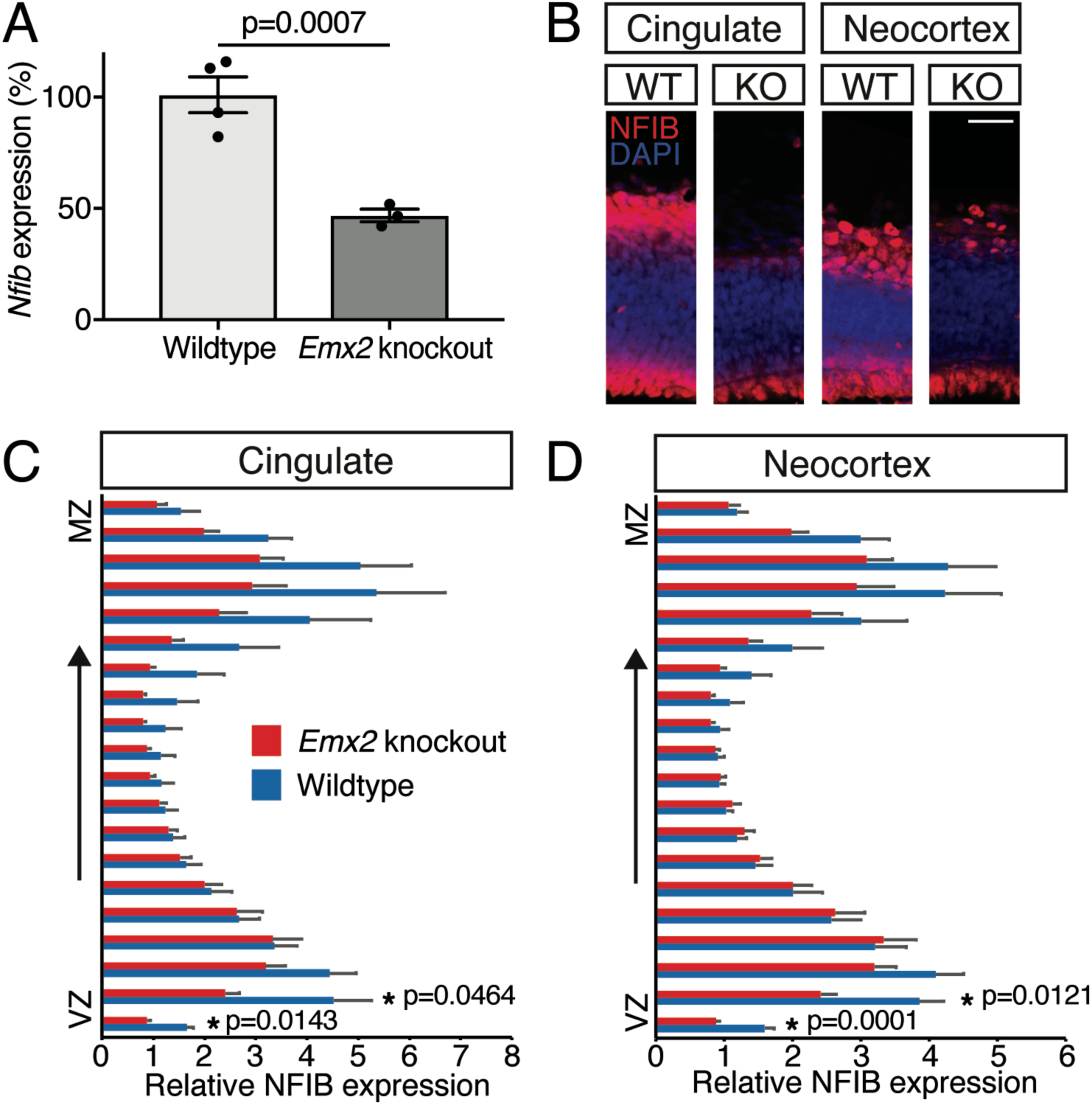
*Nfib* mRNA and protein expression is reduced in the E13.5 *Emx2* knockout cortex. (*A*) Relative *Nfib* mRNA levels in the E13.5 neocortex of wildtype (n=4) and *Emx2* knockout (n=3) littermates as determined by qPCR. Statistical significance was determined using Welch’s t-test (p = 0.0007). (*B*) Representative high-powered immunofluorescent images of the cingulate and neocortices of E13.5 wildtype and *Emx2* knockout coronal sections stained for NFIB (red) and DAPI nuclear stain (blue). Scale bar represents 50 μm. (*C*, *D*) Mean fluorescence intensity of NFIB protein expression in the cingulate cortex (*C*) and neocortex (*D*) of wildtype (blue) and *Emx2* knockout (red) sections. Fluorescence intensity was measured for each section using 20 equally spaced bins that extended from the ventricular to pial surfaces (n = 6-7 per genotype). Statistical significance was determined using unpaired t-tests.

To further test this hypothesis, we fluorescently stained matched sections of E13.5 *Emx2* knockout and wildtype littermates. The overall staining pattern of NFIB is not altered between wildtype and knockout sections, with high expression observed in the VZ and cortical plate, but not within the intermediate zone (Fig. 3B). We quantified NFIB expression within the cingulate cortex and neocortex by measuring fluorescence intensity at these regions. To normalise for variation in cortical thickness, each section was divided into 20 equal-sized bins spanning the ventricular to marginal zones (Fig 3C, D). The mean fluorescence intensity was then determined for each bin and compared between *Emx2* knockout and wildtype sections. Fluorescence intensity was significantly reduced in the VZ of *Emx2* knockout sections, but not within the intermediate zone or cortical plate. This was true for both the cingulate cortex and neocortex, where fluorescence intensity within the VZ was reduced by as much as 50% as compared to wildtype sections. Therefore, EMX2 is required for NFIB expression in radial glia.

### EMX2 transcriptionally regulates the *Nfib* promoter

We previously cloned the *Nfib* promoter region that encompasses the identified EMX2 binding sites into a luciferase reporter plasmid (Fig. 4A) (Bunt et al. 2015). To further investigate how EMX2 regulates *Nfib* expression, we tested whether luciferase activity driven via this promoter is increased in response to EMX2 over-expression. To do this, we co-transfected mouse NE-4C neuroepithelial cells or human U-251 MG glioblastoma cells with the *Nfib* promoter-driven luciferase plasmid (pNfib) and either an EMX2 over-expression or GFP control plasmid. Co-transfection of the pNfib plasmid with EMX2 increased luciferase activity by a minimum of 3-fold and 5.5-fold as compared to co-transfection with GFP in NE-4C and U-251 cells, respectively (Fig. 4B). We similarly assessed whether EMX2 over-expression had any effect on an empty luciferase reporter plasmid but did not observe any changes in luciferase activity under these conditions (control + EMX2 in Fig. 4C). These findings demonstrate that EMX2 over-expression can activate the *Nfib* promoter.

**Fig. 4.**
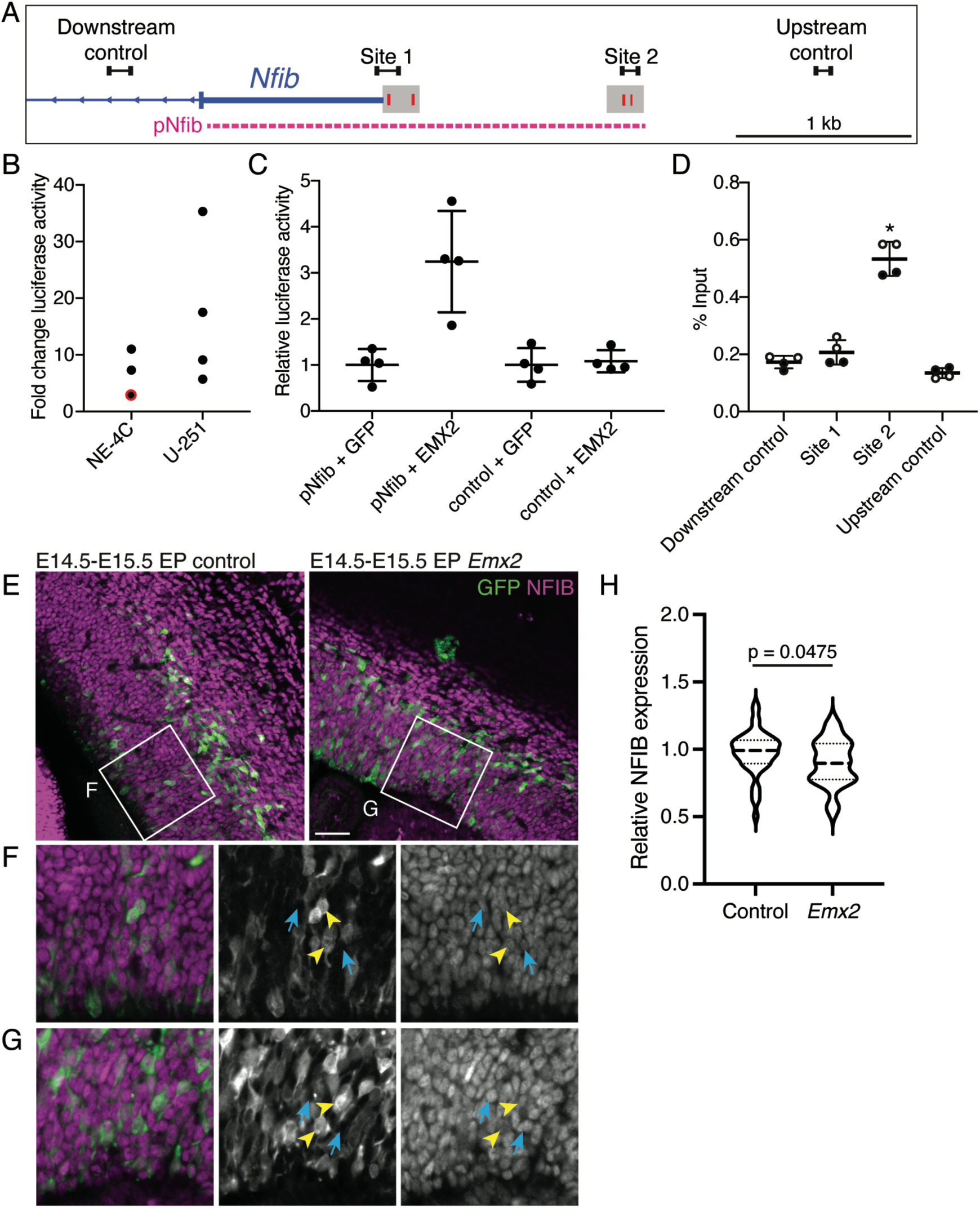
EMX2 regulates the *Nfib* promoter and fine-tunes NFIB expression. (*A*) Schematic of the proximal promoter region of *Nfib* adapted from Fig. 1. Solid blue bars represent the 5’ untranslated region and exon 1 of *Nfib*. Red boxes enclosed by grey boxes denote potential EMX2 binding sites within potential regulatory elements shared between human and mouse. The dashed line in magenta depicts the region cloned into the pNfib luciferase plasmid. Primers pairs for ChIP-qPCR are depicted by adjoining black boxes. Scale bar represents 1000 base pairs. (*B*) Relative fold change in luciferase activity measured in NE-4C and U-251 MG cells. Each data point represents an individual biological replicate, in which relative luciferase activity of the *Nfib* promoter co-transfected with EMX2 was determined upon normalisation to GFP control. Relative luciferase activity increased significantly upon EMX2 over-expression (p = 0.0466 (NE-4C) and p = 0.0068 (U-251) in ratio paired t-tests). (*C*) Representative luciferase assay performed in NE-4C cells, corresponding to the biological replicate outlined in red in (*B*). Data points represent individual technical replicates normalised to luciferase activity obtained from the respective luciferase plasmids co-transfected with GFP. (*D*) EMX2 binding (represented as % input) as assayed by ChIP-qPCR. Statistical significance was determined using unpaired t-tests. EMX2 binding was statistically significant at site 2 as compared to all other tested regions (p = 0.0219, 0.0340 and 0.0176 when compared to downstream control, site 1 and upstream control, respectively). Open and closed circles represent individual biological replicates from separate experiments (n = 2). (*E*) Representative immunofluorescent images of E15.5 wildtype coronal sections electroporated at E14.5 with control or *Emx2* plasmids. Sections were stained for NFIB (magenta) and GFP (green). Scale bar represents 50 μm. (*F*, *G*) High-powered images represented by insets in (*E*). Yellow arrowheads denote electroporated GFP-positive cells. Cyan arrows denote cells without GFP expression. (*H*) Violin plots representing relative NFIB expression in the VZ normalised to mean expression of adjacent GFP-negative cells within the same sections. Statistical significance was determined using a Mann-Whitney test. Dotted lines within individual plots denote the lower quartile, median and upper quartile (n = 7-9 per condition with 5 electroporated and 5 GFP-negative cells analysed per animal).

We next over-expressed EMX2 in NE-4C cells to determine whether the direct binding of EMX2 could be detected by ChIP-qPCR. We designed primer pairs to detect potential binding at both these regions (Sites 1 and 2 in Fig. 4A). As a control, we also designed primer pairs for regions located upstream and downstream that were void of potential EMX2 binding sites (downstream control and upstream control in Fig. 4A). We surveyed the enrichment of EMX2 binding at each of these sites following chromatin immunoprecipitation using an antibody specific for EMX2. EMX2 binding was significantly enriched at the upstream binding region within the *Nfib* promoter (Site 2 in Fig. 4A, D) as compared to all other sites surveyed (Fig. 4D). Hence, our findings demonstrate that EMX2 is capable of directly binding to the *Nfib* promoter to transcriptionally regulate its expression.

### EMX2 over-expression in the developing cortex down-regulates NFIB expression

As further proof-of-principle, we sought to determine the effect of ectopic EMX2 over-expression on NFIB expression in neural progenitor cells *in vivo*. To do this, we electroporated wildtype embryos at E14.5, an age at which endogenous EMX2 expression is decreasing, and collected embryos for analyses after 24 hours. We electroporated these embryos with a plasmid that encoded for the expression of both *Emx2* and GFP, or a control plasmid encoding GFP only. Coronal sections from both conditions were then stained with antibodies against NFIB and GFP. We compared NFIB expression between electroporated cells within the VZ of both conditions, normalised to adjacent GFP-negative cells within the same sections (Fig. 4E-G). Unexpectedly, over-expression of EMX2 resulted in a reduction in NFIB expression within the VZ where both transcription factors are co-expressed (Fig. 4G, H). Therefore, this finding suggests that besides its role in activating *Nfib* expression, EMX2 may also be capable of repressing *Nfib* in a manner dependent on its cellular context.

## Discussion

In this study, we present evidence demonstrating that EMX2 transcriptionally regulates *Nfib* during early cortical development. Using a computational approach, we identified EMX2 as a candidate transcriptional regulator of *Nfib* with conserved binding sites in human and mouse (Fig. 1A-E). This inference is supported by the positive correlation between their expression levels in published RNA-seq and scRNA-seq datasets (Fig. 1F, G). In line with their known expression patterns, immunofluorescence analyses demonstrate that both proteins are co-expressed within the VZ in similar gradients across the developing cortex (Fig. 2). Furthermore, *Emx2* deletion results in a reduction of NFIB expression within the VZ where both proteins are co-expressed but not the nascent cortical plate (Fig. 3). *In vitro* assays in NE-4C and U-251 MG cells suggest that the regulation of *Nfib* occurs through direct binding of EMX2 at the proximal promoter (Fig. 4A-D). Interestingly, ectopic over-expression of EMX2 via *in utero* electroporation does not up-regulate NFIB but leads to its repression in cells occupying the proliferative zone (Fig. 4E-H). Therefore, our findings suggest that EMX2 may function to fine-tune NFIB expression during cortical development and in this manner contribute to the timely onset of neurogenesis within this system.

We previously described a cohort of 18 individuals with *NFIB* haploinsufficiency (Schanze et al. 2018). While these individuals demonstrated variable phenotypes, the most common neurodevelopmental phenotypes observed were mild intellectual disability and macrocephaly. Analyses of *Nfib* knockout mouse models suggest that deficiencies in neuronal and glial differentiation underlie these defects (Barry et al. 2008; Piper et al. 2009; Betancourt et al. 2014; Gobius et al. 2016; Bunt et al. 2017). During cortical development, neuronal differentiation requires neural progenitor cells to switch their mode of cell division from symmetric self-renewing divisions to asymmetric neurogenic divisions. Defects in this switch could result in an expanded progenitor cell population, particularly if progenitor cells continue dividing symmetrically unhindered. Our analyses of *Nfia*; *Nfib* double homozygous knockout embryos demonstrate that the delay in neuronal differentiation observed upon loss of *Nfib* is indeed accompanied by the expansion of the neural progenitor cell pool within the proliferative VZ (Bunt et al. 2017). Transcriptome analyses performed on *Nfib* knockout embryos have also identified mis-regulated genes that may contribute to this phenotype (Betancourt et al. 2014; Bunt et al. 2017). However, our understanding of the underlying molecular pathways encompassing NFIB remains limited. Uncovering these pathways could identify additional genes involved in this process and provide novel insights into congenital disorders of cortical development.

EMX2 was previously reported as a potential rare cause of schizencephaly in humans but whether this is indeed true requires further investigation (Brunelli et al. 1996; Faiella et al. 1997; Tietjen et al. 2007; Merello et al. 2008; Hehr et al. 2010). Analyses of mouse models demonstrate that EMX2 regulates many processes throughout cortical development. Contradicting NFIB function, EMX2 expression during the initial stages of cortical development is crucial for promoting symmetric cell divisions and maintaining the progenitor cell population (Heins et al. 2001; Brancaccio et al. 2010). Loss of EMX2 during this early period results in precocious cell differentiation (Pellegrini et al. 1996; Yoshida et al. 1997; Shinozaki et al. 2002). Consequently, the progenitor cell population is rapidly depleted and less neurons are generated. Defects in neuronal differentiation are also reported to affect neuronal populations that arise from outside the cortex that do not express EMX2 (Mallamaci et al. 2000; Shinozaki et al. 2002), indicative of a non-autonomous role in the regulation of differentiation of these cell populations. Besides this, contrary to its role during early development, EMX2 expression in progenitor cells at later ages is reported to drive asymmetric divisions (Gangemi et al. 2001; Galli et al. 2002; Gangemi et al. 2006). These findings demonstrate that EMX2 functions in complex roles that require careful examination.

EMX2’s ability to regulate both symmetric and asymmetric divisions is intriguing. A potential explanation for this is that EMX2’s interaction with other transcription factors or epigenetic factors that are present in different spatiotemporal patterns alters its ability to bind to different genomic regions and regulate different target genes. Additionally, EMX2’s interaction with other transcription factors may be altered depending on the expression level of EMX2 itself. In this manner, EMX2 could potentially repress and activate NFIB expression in neural progenitor cells at different stages of development. Another potential scenario to consider is that the direct binding of EMX2 to the *Nfib* promoter elicits one response (either activation or repression of the promoter), while the opposite effect is brought about through an unknown mechanism that may be independent of EMX2 binding to the *Nfib* locus. Identifying where EMX2 binds in the genome and its target genes will be key to understanding how the function of EMX2 changes throughout development and how it is able to activate and repress NFIB in different contexts. Recent advances in the field of single-cell multiomics will be crucial for this.

To the best of our knowledge, only a few downstream targets of EMX2 have been identified in the central nervous system. These include *Wnt1* (Ligon et al. 2003), *Tenm1* (Beckmann et al. 2011) and *Sox2* (Mariani et al. 2012). Besides these, *Reln* expression is also reduced in *Emx2* knockout mice and is thought to underlie the disorganisation of radial glial fibres observed in these mice (Mallamaci et al. 2000). We previously reported a similar phenotype affecting radial glial cells in the hippocampus of *Nfib* knockout mice (Barry et al. 2008). Together with the present findings, our previous study suggest that reduced NFIB expression could be a contributing factor to the phenotype of radial glial cells in *Emx2* knockout mice.

Besides *Nfib*, the phenotypes of *Nfia* and *Nfix* knockout mice demonstrate that these genes are similarly important for cortical development (das Neves et al. 1999; Shu et al. 2003; Campbell et al. 2008; Gobius et al. 2016). Deletion of multiple *Nfi* genes results in a more severe phenotype as compared to the deletion of single *Nfi* genes, demonstrating that these transcription factors have overlapping but non-redundant functions in regulating these processes (Harris et al. 2016; Bunt et al. 2017). We postulate that the overlapping function of these transcription factors in regulating cell differentiation enables tighter control over the onset of this process during cortical development, as each gene could be individually regulated to fine-tune the total level of NFI expression. In this scenario, the precise expression of each *Nfi* gene will be important for normal development, potentially to ensure that the appropriate proportion of all major cell types are produced. Consequently, understanding how each *Nfi* gene is regulated could reveal important information regarding the regulation of cell differentiation in the developing central nervous system.

In the context of *Nfib*, EMX2 could fine-tune NFIB expression by first repressing its expression earlier in development and then activating *Nfib* expression as neurogenesis begins. This sequence of events explains why EMX2 does not activate *Nfib* prior to E11.5, but both transcription factors are expressed in similar gradients upon NFIB expression. Nevertheless, NFIB could also be repressed prior to E11.5 by other means, such as post-transcriptional regulation or the binding of other transcription factors to the *Nfib* promoter. The 3’ untranslated region (3’UTR) of *Nfib* is remarkably long at approximately 6500 nucleotides in length. The microRNA miR-153 (Tsai et al. 2014; Tsuyama et al. 2015) as well as DROSHA (Rolando et al. 2016) are capable of down-regulating *Nfib* mRNA and protein expression through binding at this region. miR-153 is particularly interesting given that its expression in the cortical VZ is detected at E9.5 but its expression in the VZ significantly decreases by E14.5 (Tsuyama et al. 2015). Besides miR-153, the miRNAs miR291-3p, miR183 and miR92 are similarly down-regulated by at least 4-fold in rat cortical progenitors when compared between E11 and E13, coinciding with the increase in *Nfib* expression in these cells (Nielsen et al. 2009). Predicted binding sites for each of these miRNAs are present within the *Nfib* 3’UTR. Hence, these miRNAs together with miR-153 could play an important role in regulating the onset of *Nfib* expression in neural progenitor cells.

While other regulators of *Nfib*, such as BRN2 (Fane et al. 2017), ASCL1 (Borromeo et al. 2016) and MYC (Mollaoglu et al. 2017) have been reported outside the central nervous system, none of these are likely to play a role in the onset of *Nfib* expression in the central nervous system given their differences in temporal and spatial expression. Nevertheless, since EMX2 expression is limited to radial glial cells throughout corticogenesis, other transcription factors are likely to play key roles in regulating *Nfib* expression, particularly in cell types that do not express EMX2 (Gulisano et al. 1996; Mallamaci et al. 1998) and in the adult brain where NFIB but not EMX2 is widely expressed (Gangemi et al. 2001; Chen et al. 2017). EMX1 is a fitting candidate given its expression in both progenitor cells and postmitotic neurons during cortical development (Gulisano et al. 1996). *In vitro* experiments demonstrate that EMX1 and EMX2 share identical consensus binding motifs (Jolma et al. 2013). Therefore, EMX1 could potentially bind to the EMX2 binding sites that we identified within the *Nfib* promoter to similarly regulate *Nfib* expression in progenitor cells and postmitotic neurons.

Another scenario that should be considered is the prospect of the NFI transcription factors, including NFIB itself, potentially regulating *Nfib* expression. A super enhancer was recently discovered within the intragenic region adjacent to the *Nfib* locus that is active in neural stem cells. This super enhancer was identified through the enrichment of the Mediator complex within this region, and is also bound by the transcription factor TCF4 in neural stem cells (Moen et al. 2017; Quevedo et al. 2019). Chromatin immunoprecipitation performed with a pan-NFI antibody (recognising all four NFI transcription factors) similarly demonstrates NFI binding at this region (Mateo et al. 2015). Since NFIB is capable of directly interacting with TCF4 as well as MED15 (a subunit of the Mediator complex) (Moen et al. 2017; Quevedo et al. 2019), the onset of *Nfib* expression followed by its potential binding at this region could result in a feed-forward loop that sustains NFIB expression in the central nervous system. Hence, besides EMX2, NFIB itself could be a potential regulator of *Nfib* expression during cortical development.

In conclusion, this study demonstrates that EMX2 regulates *Nfib* expression in cortical progenitor cells that occupy the dorsal VZ of the developing telencephalon. Given its earlier onset and similar expression patterns when compared to NFIB, EMX2 could be important for driving the initiation of *Nfib* expression during cortical development but may also function to repress *Nfib* expression depending on context. In this way, EMX2 could fine-tune NFIB expression to regulate the onset of cell differentiation during development.

## Acknowledgements

We thank the staff of the University of Queensland Biological Resources (UQBR) animal facility and the QBI Advanced Microimaging and Analysis Facility for their expertise and support in this project. We thank Peter Gruss (Max Planck Society, Munich, Germany) for providing the *Emx2* knockout mouse line and Magdalena Götz for providing the pMXIG-Emx2 plasmid. We are also grateful to Yunan Ye and Elizabeth Vanderkop for their assistance with experiments.

## Funding

This work was supported by the National Health and Medical Research Council (GNT1100443 and GNT1120615) and the Australian Research Council (DP150104748 and DP200102363). Students were supported by the University of Queensland (UQ Centennial Scholarship to JWCL and International Postgraduate Student Scholarship to K-SC) and the Australian Government (Research Training Program Scholarship to JWCL).

## Author contributions

Conceptualization: JWCL, JB, LJR; Formal analysis and investigation: JWCL, JB, CRB, CM, MTJ, K-SC; Writing - original draft preparation: JWCL, JB, LJR; Writing - review and editing: JWCL, JB, CRB, CM, MTJ, K-SC, LJR; Resources: RMG; Supervision: JWCL, JB, LJR

## Compliance with Ethical Standards

## Conflicts of interest

All authors declare that they have no conflicts of interest.

## Ethical Approval

All applicable international, national, and/or institutional guidelines for the care and use of animals were followed. This article does not contain any studies with human participants performed by any of the authors.

## Notes

### Competing Interest Statement

The authors have declared no competing interest.

