## Supplementary Figure for "EMX2 transcriptionally regulates *Nfib* expression in neural progenitor cells during early cortical development"

**Supplementary material**

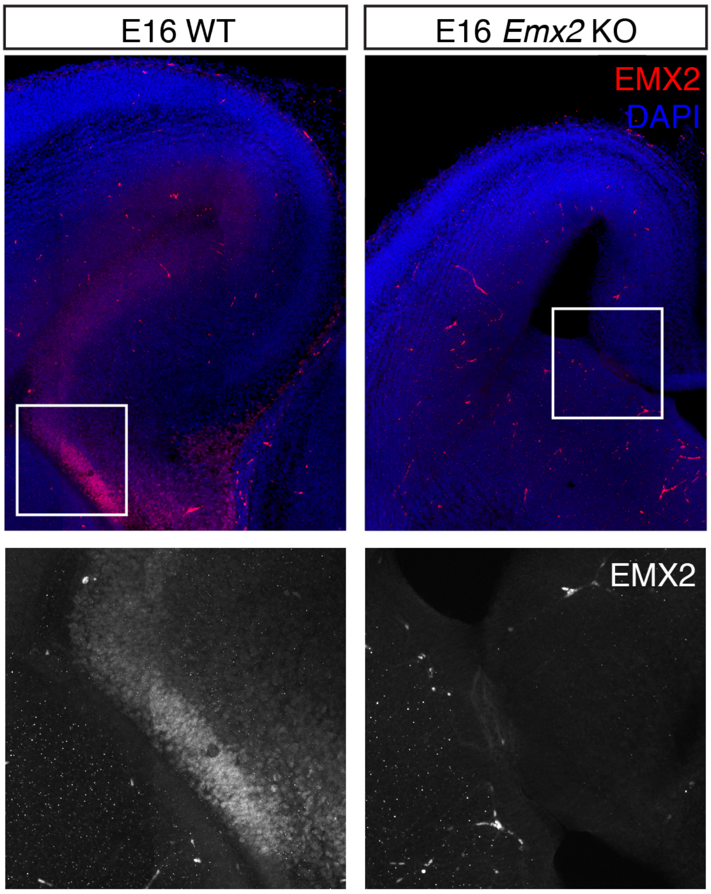


**Supplementary Fig. 1**

Validation of specificity of the rabbit anti-EMX2 (HPA065294, Atlas Antibodies) antibody used for immunofluorescence analyses. E16 wildtype (WT) and *Emx2* knockout sections were labelled for EMX2 (red) and DAPI nuclear stain (blue). EMX2 staining is observed in wildtype but not in knockout sections.
